# Seed sourcing for climate-resilient grasslands: the role of seed source diversity during early restoration establishment

**DOI:** 10.1101/2021.12.15.472808

**Authors:** Jessica Lindstrom, Marissa Ahlering, Jill Hamilton

## Abstract

Restoration often advocates for the use of local seed in restoration, however increasingly new strategies have been proposed to incorporate diverse sources to maintain evolutionary potential within seed mixes. Increasing seed sources per species within a seed mix should increase genetic variation, however, few empirical studies have evaluated how seed source diversity impacts plant community composition following restoration. Thus, the goal of this research was to compare the use of single or multi-source seed mix treatments to plant community diversity following restoration. Using 14 species commonly applied in grassland restoration, we examined plant community diversity following restoration comparing seed mixes with either one or five sources per species across two restoration sites in Minnesota and South Dakota, United States. Following seeding, species establishment and abundance were recorded to calculate plant diversity for each seed mix treatment. There were no major effects of seed mix treatment on community emergence and diversity observed, with the majority of plant establishment reflecting non-seeded species. However, site-specific differences were observed. Heterogeneous land-use history associated with the Minnesota site likely contributed to differences across the restoration treatments. In contrast, community diversity at the South Dakota site was homogeneous across seed mix treatments with changes in plant community influenced solely by early season species establishment. This suggests land-use history irrespective of seed mix treatment influences establishment and persistence, particularly in the first year following restoration. Future monitoring across seasons will be needed to evaluate if community diversity changes in response to seed mix treatment.

## INTRODUCTION

One of the major aims of ecological restoration is to restore or re-establish functional plant species diversity to ensure key ecosystem services are maintained (Barr et al. 2017; Montoya et al. 2012). To ensure ecosystem health and the maintenance of productive plant communities, this includes creating diverse seed mixes for application in restoration (Tilman et al. 1996, 1997, 2001; Brudvig 2011). These seed mixes create communities that may be resilient to changes in nutrient availability (Craven et al. 2016), competition from non-natives (Funk et al. 2008; Oakley & Knox 2013; Yurkonis et al. 2012; Norland et al. 2013), and climate change (Isbell et al. 2015). Evolutionary theory emphasizes the important role both inter- and intraspecific variation established within seed mixes may have to restoration success over time (McKay et al. 2005). Greater biodiversity within restoration communities may increase total plant productivity across time leading to increased stability in soil nutrient availability (Craven et al. 2016), and resilience to extreme events (Isbell et al. 2015). In addition, intraspecific variation is essential as this may provide the raw material that natural selection may act upon and is needed to maintain species’ evolutionary potential (Pizza et al. 2021; McKay et al. 2005). Despite the importance of intraspecific diversity to restoration success, few studies have quantified the role diversity within species has to restoration outcomes (Hamilton et al. 2020). Consequently, to ensure that plant communities persist over time and in response to change, there is a need to consider both the role of within and between species diversity to restoration.

Current strategies used to establish seed mixes often advocate a ‘local is best’ approach (Broadhurst et al. 2008; McKay et al. 2005). This approach assumes that local seed sources will have greatest fitness in local restoration environments relative to non-local sources (Kawecki & Ebert 2004; Hoban et al. 2016). While there is evidence of local adaptation for many plant species (Leimu et al. 2010; Hereford 2009), the degree or scale of adaptation is often unknown (McKay et al. 2005). Furthermore, to conserve evolutionary potential requires genetic variation (Kawecki & Ebert 2004). Genetic diversity is the raw material that selection acts upon and is necessary for adaptation to changing environmental conditions. Genetic variation may be lost through random fluctuations in population size via genetic drift, but maintained through gene flow among populations (Reed & Frankham 2003). In addition, small, isolated plant populations that exhibit reduced connectivity or gene flow may exhibit reduced genetic variation, but increased genetic differentiation (Durka et al. 2017). If seeds are sourced locally for restoration from small, isolated populations then individual seed sources may not have the requisite genetic variation needed to adapt to change (Davis et al. 2005; Etterson & Shaw 2001). To ensure the maintenance of evolutionary potential therefore may require seed sourcing strategies that increase genetic diversity. Accounting for the role evolutionary forces play in the maintenance of diversity will aid in establishing seed mixes that ultimately increase restoration success (Bucharova et al. 2017; Hamilton et al. 2020).

To ensure preservation of evolutionary potential, variation within species is required alongside the establishment of species rich seed mixes. The combination of intraspecific and interspecific species diversity can influence community composition during establishment (Larson et al. 2013). Diversity at these two scales can impact short-term response to the environment and competition with local seed banks (Grman et al. 2013). During the first few years following restoration it is expected that communities will be largely dominated by non-seeded weedy species typically found within the soil seed bank (Bakker et al. 1996). For example, when comparing an active prairie restoration to multiple remnant prairies, Martin et al. (2005) observed more non-native species present within the restoration, with the overall proportion of non-natives ranging from 236% to 413% higher in the restoration relative to remnant sites. Thus, considering early establishment of seeded relative to non-seeded species may be important to predicting longer-term plant community composition. Despite the potential importance of early establishment to long-term restoration success, this phase is often overlooked in favor of evaluating restorations after they have been established for several years.

Globally, native grasslands remain one of the most critically imperiled ecosystems requiring active restoration (Hoekstra et al., 2005). These ecosystems provide essential services, including maintenance of hydrological flow and retention (Seeling & DeKeyser 2006), carbon sequestration (Euliss et al. 2006), nutrient cycling, and habitat for a diversity of species (Helzer & Jelinski 1999; Skagen et al. 2008). Throughout the North American Great Plains, up to 87% of historical grassland habitat has been lost primarily to agricultural conversion (Comer et al. 2018; Hoekstra et al. 2005; Samson et al. 1999) leading to highly fragmented and isolated remnant habitats. Where these grasslands remain, they are prone to invasion by non-native species and the evolutionary consequences of isolation, which has lasting negative effects to diversity and species richness (DiAllesandro et al. 2013; Haddad et al. 2015). Ensuring seed mixes restore grassland populations so they have the capacity to adapt to change, resist invasion, and persist over time is critical. However, the role of intraspecific diversity within seed mixes to restoration success has yet to be empirically evaluated. Therefore, it is necessary to consider the impact of both species and population diversity within seed mixes has to establishment of grassland restorations.

We assessed plant community diversity following restoration using single- and multi-source seed mixes to test the role within-species seed source diversity played in community establishment. We used seed collected from five unique populations for each of 14 different species as a proxy for creating genetic diversity within a seed mix. We expected that increasing the number of unique seed sources per species used within a seed mix would lead to increased emergence diversity following restoration relative to the use of a single seed source seed mix (Bucharova et al. 2018). Overall, we predicted greater within-species diversity for seed mixes would lead to increased species diversity in restored plant communities. This research empirically evaluates the role of within species to between species diversity following restoration. This study will provide a baseline understanding of the role of diversity across scales to establishment during restoration.

## METHODS

### Seed Collection

In the summer of 2019, seed from 12 forb and two grass species were collected between June and October from remnant native prairies within the Northern Great Plains of the United States. A minimum of five unique populations per species each were collected from the Missouri Coteau region of North and South Dakota and from the northwestern prairie region of Minnesota (Table 1, Fig. 1). These 14 species were chosen because they are widely distributed throughout the Northern Great Plains and are commonly used in regional restoration seed mixes (e.g., Smith 2010; Kurtz 2013). In addition, to control for potential dominance of warm-season grasses and to increase establishment of sown forbs, species chosen were weighted toward forb species (McCain et al. 2010; Norland et al. 2013; Dickson & Busby 2009). Populations were classified as distinct if separated by at least one mile, however, were more commonly spaced further apart. In northwestern MN, distances between seed source locations ranged from 3 km to 215 km (Table S2), and pairwise distances between the restoration site and seed source ranged from 2 km to 129 km (Table S3). Within the Missouri Coteau region, distances between seed source locations ranged between 2 km and 312 km (Table S4), and pairwise distances between the restoration site and seed source site ranged from 3.5 km to 214 km (Table S5).

**Table 1.** Species used experimental restoration plots for RSC and ORD sites, weighed amounts used in individual seed mix treatments, individual species composition within seed mixes, approximate seeds/m^2^, and seed viability included where applicable.

| Species Scientific Name | RSC |  |  |  |  | ORD |  |  |  |
| --- | --- | --- | --- | --- | --- | --- | --- | --- | --- |
|  | Single-<br>population<br>Seed Mix (g) | Five-<br>population<br>Seed Mix (g) | Species<br>composition<br>in mix (%) | Seeds/m<sup>2</sup> | Viable<br>Seed<br>(%) | Single-<br>population<br>Seed Mix (g) | Five-<br>population<br>Seed Mix (g) | Species<br>composition<br>in mix (%) | Seeds/m<sup>2</sup> |
| *Amorpha canescens* | 12.5 | 2.5 | 21.8 | 784 | 20 | 12.5 | 2.5 | 23.2 | 784 |
| *Anemone cylindrica* | 7.5 | 1.5 | 13.1 | 764 | 82 | - | - | - | - |
| *Artemisia frigida* | 0.5 | 0.1 | 0.9 | 556 | 62 | - | - | - | - |
| *Bouteloua curtipendula* | 5.0 | 1.0 | 8.7 | 233 | - | 1.5 | 0.3 | 2.8 | 70 |
| *Bouteloua gracilis* | - | - | - | - | - | 0.5 | 0.1 | 0.9 | 78 |
| *Dalea purpurea* | 5.0 | 1.0 | 8.7 | 353 | - | 5.0 | 1.0 | 9.3 | 353 |
| *Echinacea angustifolia* | 8.0 | 1.6 | 13.9 | 219 | - | 8.0 | 1.6 | 14.8 | 219 |
| *Geum triflorum* | 1.3 | 0.3 | 2.2 | 132 | 47 | 8.5 | 1.7 | 15.8 | 899 |
| *Helianthus maximiliani* | 1.3 | 0.3 | 2.2 | 64 | - | 1.0 | 0.2 | 1.9 | 51 |
| *Helianthus pauciflorus* | 2.0 | 0.4 | 3.5 | 31 | - | 2.0 | 0.4 | 3.7 | 31 |
| *Hesperostipa comata* | - | - | 1.5 | - | - | 0.6 | 0.1 | 1.2 | 18 |
| *Liatris punctata* | 4.3 | 0.9 | 7.4 | 117 | - | 1.0 | 0.2 | 6.5 | 274 |
| *Pedimelum argophyllum* | 2.5 | 0.5 | 4.4 | 88 | - | 3.5 | 0.7 | 1.9 | 27 |
| *Potentilla arguta* | 1.5 | 0.3 | - | 1352 | 88 | 1.3 | 0.3 | 6.5 | 123 |
| *Ratibida columnifera* | - | - | 2.6 | - | - | 2.5 | 0.5 | 2.3 | 1127 |
| *Schizachyrium scoparium* | 2.8 | 0.6 | 4.8 | 162 | - | - | - | 4.6 | 412 |
| *Solidago rigida* | 2.5 | 0.5 | 4.4 | 402 | 54 | 2.5 | 0.5 | 4.6 | 402 |

**Fig. 1.**
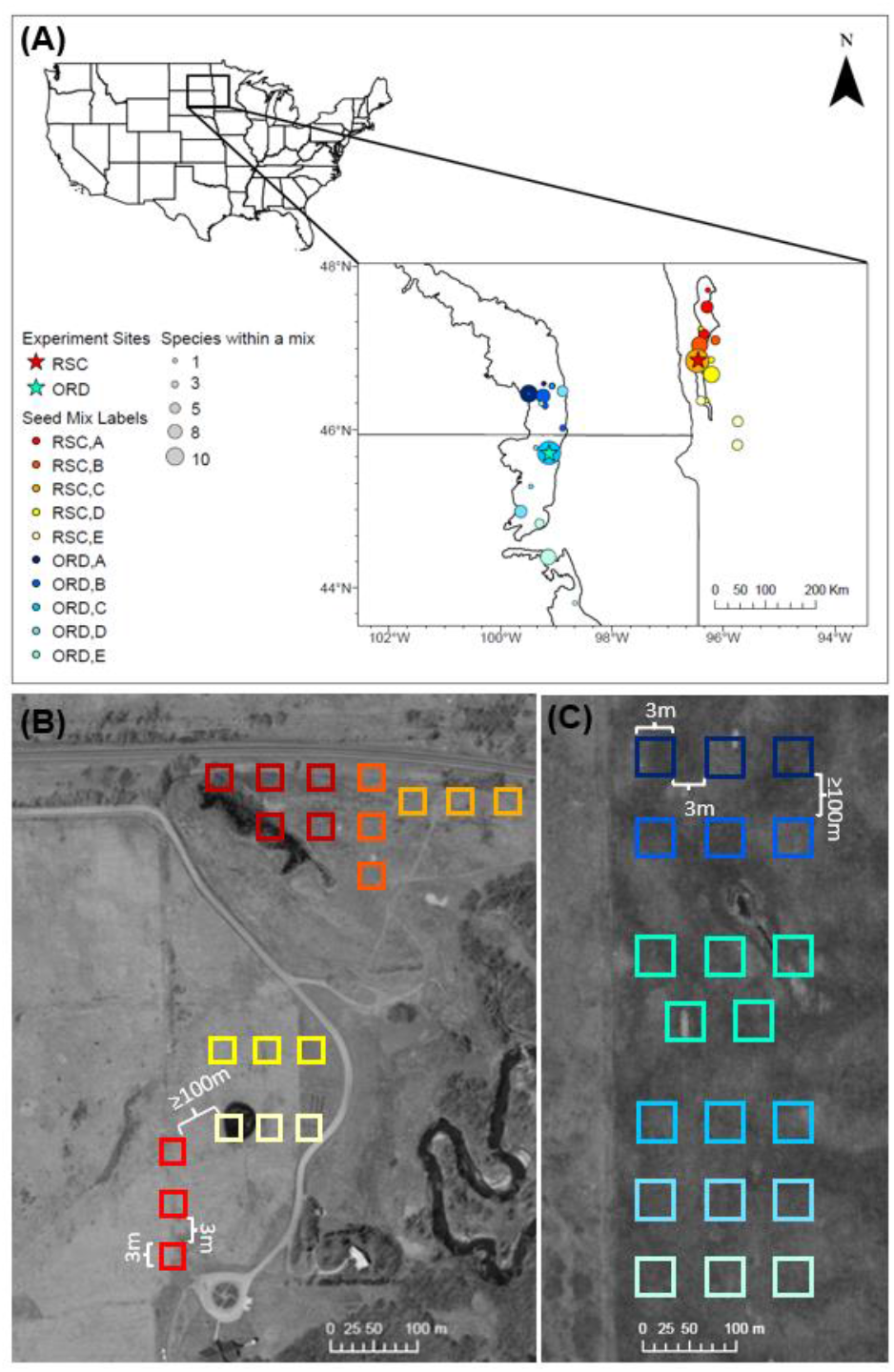
A) Seed collection sites for seed mix treatments for Missouri Coteau (blues) and northwestern MN (reds) regions respectively. Colors represent individual seed mixes, and proportional symbols indicate the number of species sourced from a single site used within a seed mix. Stars indicate experimental site locations. B) Seed mix treatment layout at the RSC restoration site in Glyndon, MN. C) Seed mix treatment layout at ORD restoration site in Leola, SD. Colors correspond to seed treatment, single-source treatments include three replicate plots and the multiple-source treatment includes five replicate plots.

Seed was hand-harvested as it ripened, with seed harvested multiple times at different sites throughout the growing season following Bureau of Land Management seed harvesting guidelines (BLM 2015). Within each population, individual maternal seed heads were sampled at least three feet apart to reduce potential relatedness within populations. For species with multiple seed heads, no more than 30% of available seed per maternal seed head was collected.

### Seed Mix Preparation

Following harvest, seeds were dried at room temperature for a minimum of two weeks and then transferred to 4°C storage for seven months to provide cold stratification and maintain viability. Seeds were cleaned using several species-specific approaches. Large seeds were stripped by hand, smaller seeds separated using sieves, *Hesperostipa comata* (Needle and thread grass) seed awns were trimmed during the drying process to limit tangling, and *Solidago rigida* (Stiff Goldenrod) and *Helianthus maximiliani* (Maximilian sunflower) and *H. pauciflorus* (Stiff sunflower) seed were mechanically cleaned and separated using a Fractioning Aspirator Test Model at the USDA Agricultural Research Center in Fargo, ND.

Seed was weighed for each species (Mettler Toledo, ML503T/00) from each population to calculate population-specific numbers of seeds using a seeds per gram conversion (Table S1). To maximize the seeds per species in the mix and ensure seed mix consistency across treatments and replicates, the amount of seeds to include in the mix per species was calculated based on the population with the lowest seed weight (g). In addition, for *Artemisia fringida* (Fringed sagewort), *H. pauciflorus*, and *S. rigida*, the amount of seeds used in the seed mixes was reduced by 0.9%, 3.5-7.0%, and 4.4-6.0% of the lowest seed weight respectively, to ensure these species were not overrepresented in seed mixes as they can exhibit dominant characteristics (Table 1).

Across the two regions, seed mixes were established using the same species with the exceptions of *A. fringida, Anemone cylindrica* (Tall timbleweed), and *Schizachyrium scoparium* (Little bluestem), which were collected and planted exclusively in the northwestern MN region and *Ratibida columnifera* (Prairie coneflower), *H. comata, Bouteloua gracilis* (Blue grama), which were collected and planted exclusively in the Missouri Coteau region. Five different seed mixes were established each using a single unique population per species for the seed mix within each of the two restoration regions. For these single-source seed mixes, populations for the different species were largely sourced from similar latitudes to minimize potential impacts associated with latitudinal variation in phenology (Olsson & Ågren 2002; Dunnell & Travers 2011) (Fig. 1). In addition to five single source seed mixes, one multiple-source seed mix was established for each region. The multi-source seed mix used proportionally the same amount of seeds per species as the single-source mix, but each species’ contribution was divided evenly across five population sources. Thus, for both single and multi-source seed mixes the proportion of seed used per species was the same. In this way, the ratio of species present within the single source and multi-source was maintained across seed mixes for direct comparison. Vermiculite (Vigoro) was added to final seed mixes in a 1:1 ratio as a common method to increase seed to soil contact during planting and thus increase probability of emergence (Shaw et al. 2020).

### Seed Viability

Unused seed from the restoration plots sampled from the northwestern MN region were sent to South Dakota State University’s Seed Testing Laboratory to assess seed viability. Unused seed from the Missouri Coteau were not available for seed viability testing. These tests evaluated the total viability of individual species when grown under ideal laboratory growth conditions to induce germination. This test reported the percent of seed that germinated defined as the total number of individuals emerged per seeds planted, percent of hard seed defined as seed that is dormant due to a water impervious seedcoat, and dormant seed which is defined as seed that is viable but does not germinate due to a physical or physiological condition (SDSU Seed Testing Laboratory; https://www.sdstate.edu/sites/default/files/file-archive/2021-07/Seed-Testing-Lab.pdf).

### Restoration Sites and Site Preparation

During May and June of 2019, experimental restoration sites were identified and prepped in both the northwestern MN and Missouri Coteau regions. The northwestern MN restoration site was established at the Minnesota State University Moorhead Regional Science Center (RSC) (46.872, -96.452) in Glyndon, MN. Portions of this site are abandoned agricultural brome fields that are adjacent to remnant mesic prairie owned by Buffalo River State Park. Another portion of this site was actively maintained as the Ponderosa golf course starting in 1962 and continued operation after the transfer of ownership until May 2015, following which limited mowing management has occurred. Due to site and space limitations, both areas of this site were used to establish the experimental plots. The Missouri Coteau restoration site was established on the Samuel H. Ordway Prairie Preserve (ORD) (45.704 -99.086), owned and managed by The Nature Conservancy (TNC). Prior to TNC ownership in 1978, this site was used as a brome/alfalfa production plot for cattle. Since TNC’s ownership, this site has been maintained for hay production every other year.

In 2019, the RSC site was prepared by placing landscape cloth over experimental restoration plots to remove existing vegetation and limit potential establishment and competition with the existing seedbed prior to applying the restoration treatment. In fall 2019, the ORD site was treated with herbicide prior to application of restoration treatment (Roundup®, 3-4% concentration) within each plot to reduce competition with existing weedy vegetation during establishment. Additionally, all plots had a second Roundup treatment in early May, 2020 to further reduce *Bromus inermis* (Smooth brome) encroachment.

At each site, twenty 3 × 3m experimental restoration plots were established. This included establishment of five different single source seed treatment plots each replicated three times (n=15) and one multi-seed source treatment replicated five times (n=5). For each individual replicated plot within a seed treatment, a barrier of 3m was maintained and a minimum 100m buffer maintained between each single- and multi-source seed treatment group to limit potential gene flow between plots.

### Planting Experimental Restoration Treatments

To establish the restoration treatments, tarps were removed from the plots at the RSC site, and litter was raked and hand weeded in April 2020 at both sites to expose the seed bed. Following this, each plot was broadcast seeded and then raked again to increase seed-soil contact. For both sites, five times the total commonly recommended seeding rate of ∼5kg (11 pounds) of seeds per acre were applied to increase probability of emergence success (Rowe 2010). Higher seeding rates were applied as these rates have previously been associated with increased establishment and diversity following restoration (Sheley & Half 2006; Barr et al. 2017). An agri-fab push lawn roller was used to increase seed to soil contact and enhance the probability of germination success. To limit potential carryover of seeds between seed treatments the roller was rinsed and dried between each application. Finally, each plot received a one-time watering treatment. Throughout the growing season, plot maintenance included weekly barrier mowing around each plot. In July, mid-season mowing was performed at both sites to increase light availability and reduce competition with non-seeded species (Maron & Jefferies 2001; Kaul & Wilsey 2020). Plots were mowed at the maximum adjustable height setting (12.7cm) and all trimmings were removed.

### Data Collection

Each restoration plot was visited once per month at both sites between June and September of 2020 to assess plant community composition. A 0.2m x 0.2m quadrat randomly placed at each of the four cardinal corners and center of each replicated experimental plot was used to estimate community composition of the broader restoration plot. To reduce the impact of edge effects, quadrats were not placed directly at the edges of each plot. For all species present in the quadrat, we counted the number of individuals present and estimated the percent cover per species. Individuals that were unidentifiable in the field were marked with unique toothpicks and photographed for later identification. There were two unknown species at the RSC site and three at the ORD site that did not match planted species seedlings and were unable to be identified.

These species were uniquely labeled as unknowns and included in diversity calculations as unique non-seeded species. Total percent cover of dead vegetation and percent bare soil cover was also assessed visually within the quadrat. At the quadrat-level, total species coverage was recorded as the total percent coverage of each species, litter coverage was the percent cover of dead matter covering the ground, and soil coverage was the percent of visible bare ground. Each coverage estimate was assessed with a modified Daubenmire cover-class system for grassland vegetation (Table S6; Daubenmire 1959) and averaged across quadrats to obtain replicate-level percent coverages for each seed mix treatment.

### Statistical Analysis

To infer plant productivity and assess plant community composition following restoration, species diversity metrics such as richness, evenness, abundance, and associated diversity indices are often used and may be monitored over time (Martin et al. 2005; Polley et al. 2003). We tested for differences in community composition based on seed mix treatment at each of our restoration sites using measures of species richness and diversity. Species richness was defined as the total number of species present across all five quadrats sampled per replicate and abundance as the total number of individuals present per species across quadrats. We also analyzed the total number of unique species and the number of seeded species that established within seed treatments for replicated plots. To evaluate our seed treatment communities regardless of planted or non-seeded species status, we calculated Shannon’s Diversity Index (H’) for each seed treatment and each replicate plot across time from June to September.

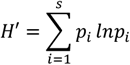

Where *s* is the total number of species within the community (richness), *p*_*i*_ is the proportion of each species (*i*) within the community relative to the total number of species multiplied by the natural logarithm and summed across all species to get a value between 0-1. Values closer to 0 indicated lower diversity and values closer to 1 indicated higher diversity. We used Shannon’s Diversity for our data as it was the most appropriate given our data collection approach (Magurran 2004). Diversity indices were calculated at the seed treatment level and for the individual replicates within seed treatment to create distance matrices.

To compare plant community diversity within each restoration site for varying seed treatments across time we used non-metric multidimensional scaling (NMDS) in three dimensions with a Bray—Curtis dissimilarity distance matrix which was derived from the Shannon’s diversity indices for each seed treatment and each month of data collection. We used NMDS because it uses an ordination approach where community data is summarized on two-axes and communities that are more similar cluster together (Ruiz-Jaen & Aide 2005).

To evaluate differences between community compositions, we performed permutational ANOVAs (PERMANOVAS) on the same Shannon’s diversity values for seed treatment communities across each month, using the adonis function in package “vegan” (Oksanen et al. 2020). We used a PERMANOVA approach to evaluate differences between individual seed treatments and more broadly between single source and five-source community diversity. Seed treatment, replicate, month, and the interaction of seed treatment and month were predictor variables and the percent bare ground and thatch were included as random-effect variables within our models. Post-hoc pairwise comparisons were performed to evaluate differences between seed mix treatment per month for RSC communities and by month for ORD communities within the pairwise.adonis function in package “pairwiseAdonis” (Martinez Arbizu 2019). All analyses were conducted in R version 4.0.2 (R Core Team 2016).

## RESULTS

### Seed Viability

Six of the 14 species sent for testing had enough seed for an assessment of viability. Variability in seed viability may impact how individual species may or may not establish within the first year following restoration. Seeds from *Amorpha canescens* exhibited a viability score of 20% with 16% of seed reaching germination, 4% labeled as hard seed, and 0% dormant seed. Seeds from *Anemone cylindrica* exhibited 82% viability with 75% of seed reaching germination, 0% hard seed and 7% assessed as dormant. Seeds from *Artemisia fringida* were 62% viable, with 25% of seed reaching germination, 0% labeled as hard seed, and 37% dormant seed. *Geum triflorum* seed had a total viability of 47% with 47% of seed reaching germination, 0% labeled as hard seed, and 0% dormant seed. *Potentilla arguta* seed exhibited 88% viability, with 66% of seed reaching germination and 0% labeled as hard seed, 22% dormant seed. Finally, *Solidago rigida* seeds had a viability score of 54% with 44% of seed reaching germination, 10% labeled as hard seed, and 10% dormant seed.

### Plant Community Structure following Restoration

Seed mix application at both the RSC and ORD sites resulted in a mixture of seeded and non-seeded species emergence. At the RSC site, seeded species emerged from all plots excluding seed treatment ‘D’ in the first growing season. Of seed mix treatment types, the multi-source seed mix type ‘ABCDE’ had the greatest number of seeded species emerge, including *Echinacea angustifolia, Helianthus maximilani*, and *Verbena hastata*. Across all seed treatments at the RSC site, *Helianthus maximilani* exhibited the greatest rate of emergence, followed by *Liatris punctata*. In the first year of observation, in total only five of the seeded species established at the RSC site. At the ORD site, seeded species emerged within all plots in the first growing season. Of seed mix treatment types, the multiple-source seed mix type ‘ABCDE’ and the single-source seed treatment ‘C’ had the greatest number of seeded species emerge, including *H. maximiliani, S*.*rigida* which were found within every seed treatment, followed by *Ratibida columnifera*, and *Dalea purpurea*. In total only six unique seeded species established at the ORD site.

At both restoration sites, seed treatment plots were largely dominated by non-seeded species (Fig. 2.) At the RSC site the most common species within our experimental restoration plots were *Ambrosia psilostachya* (Western Ragweed), *Melilotus sp*. (Sweetclover sp.), *Panicum capillare* (Witchgrass), *Poa pratensis* (Kentucky Bluegrass), *Oxalis stricta* (Yellow Wood Sorrel), *Trifolium repens* (White Clover). At the ORD site the most common species within our experimental restoration plots were *A. absinthium, Bromus inermis* (Smooth Brome), and *P. pratensis*.

**Fig. 2.**
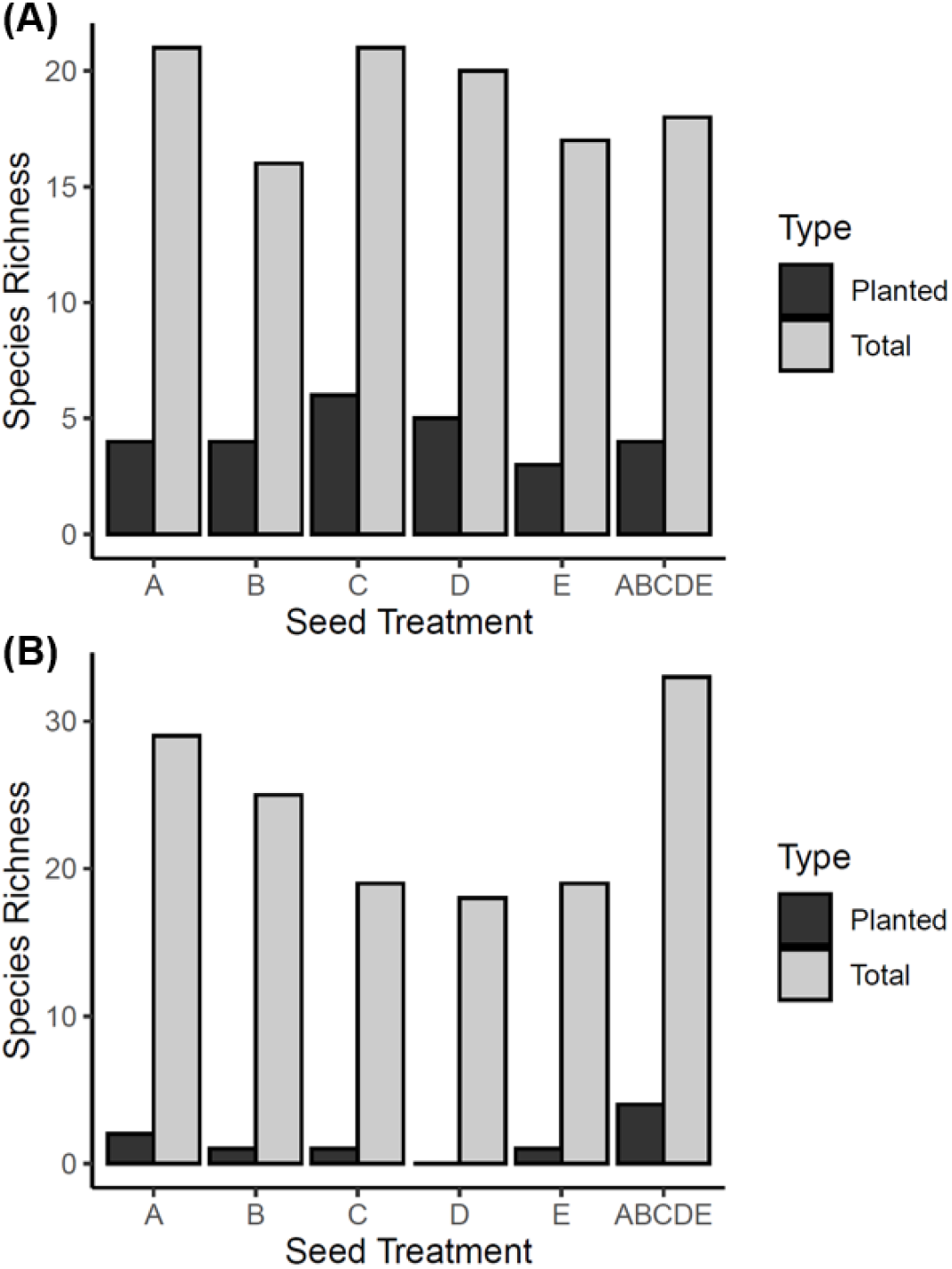
Comparison of planted and total species richness within each seed mix treatment throughout June-September 2020 within the ORD experimental site (A) and RSC experimental site (B). Overall planted richness was greater in within all ORD seed treatments compared to RSC. Total species richness was higher in RSC than in ORD, and the multiple-source seed treatment, labeled ABCDE, had greatest planted species richness.

To evaluate plant community-level differences between seed mix treatment types and across the growing season, we used a PERMANOVA based on Shannon’s Diversity. Additionally, to visualize any differences in these plant communities we used an NMDS with Bray–Curtis dissimilarity. Within the RSC site, we found significant community-level differences based on seed mix treatments (pseudo-*F*= 18.268, *p* = 0.001;), plot replicate (pseudo-*F* =7.868, *p* = 0.001), month (pseudo-*F*= 2.677, *p* = 0.018), and the interaction of seed treatment and month (*Pseudo-F*= 2.172, *p* = 0.008; Table 2). However, as very few seeded species established across seed mix treatments, the differences observed appear to be largely driven by site-level differences associated with spatial heterogeneity in the presence of non-seeded species (Fig. 3B). To then evaluate which seed treatments, or location of seed treatments within the RSC site were compositionally different, we then performed individual pairwise analyses. Pairwise comparisons evaluating community compositions differences across seed mix treatments were subset by month to account for the significant interaction of seed treatment and month found within our PERMANOVA results. From these comparisons we found the five-source seed treatment was significantly different from all single-source seed treatments across all months with the sole exception of seed source ‘E, which became more similar to the five-source treatment over time (Table S7). This follows our expectation that the multiple-source treatment would produce a more diverse community when compared to single-source treatments; however, with the caveat that differences observed seem to be driven largely by the diversity of non-seeded species present within individual plots.

**Table 2.** PERMANOVA results for community composition differences within RSC experimental seed mix treatments, using Seed Treatment, Plot Replicate, Month, and the interaction between seed treatment and month as main explanatory variables.

|  | Df | SS | MS | Pseudo F | R2 | *P* |
| --- | --- | --- | --- | --- | --- | --- |
| Seed Treatment | 5 | 1.475 | 0.295 | 18.268 | 0.323 | *0.001* |
| Plot Replicate | 14 | 1.778 | 0.127 | 7.868 | 0.390 | *0.001* |
| Month | 3 | 0.130 | 0.043 | 2.677 | 0.028 | *0.018* |
| Bare Ground | 1 | 0.002 | 0.002 | 0.135 | 0.000 | 0.897 |
| Thatch | 1 | 0.007 | 0.007 | 0.408 | 0.001 | 0.640 |
| Treatment:Month | 15 | 0.526 | 0.035 | 2.172 | 0.115 | *0.008* |
| Residuals | 40 | 0.646 | 0.016 |  | 0.142 |  |
| Total | 79 | 4.5627 |  |  | 1 |  |

**Fig. 3.**
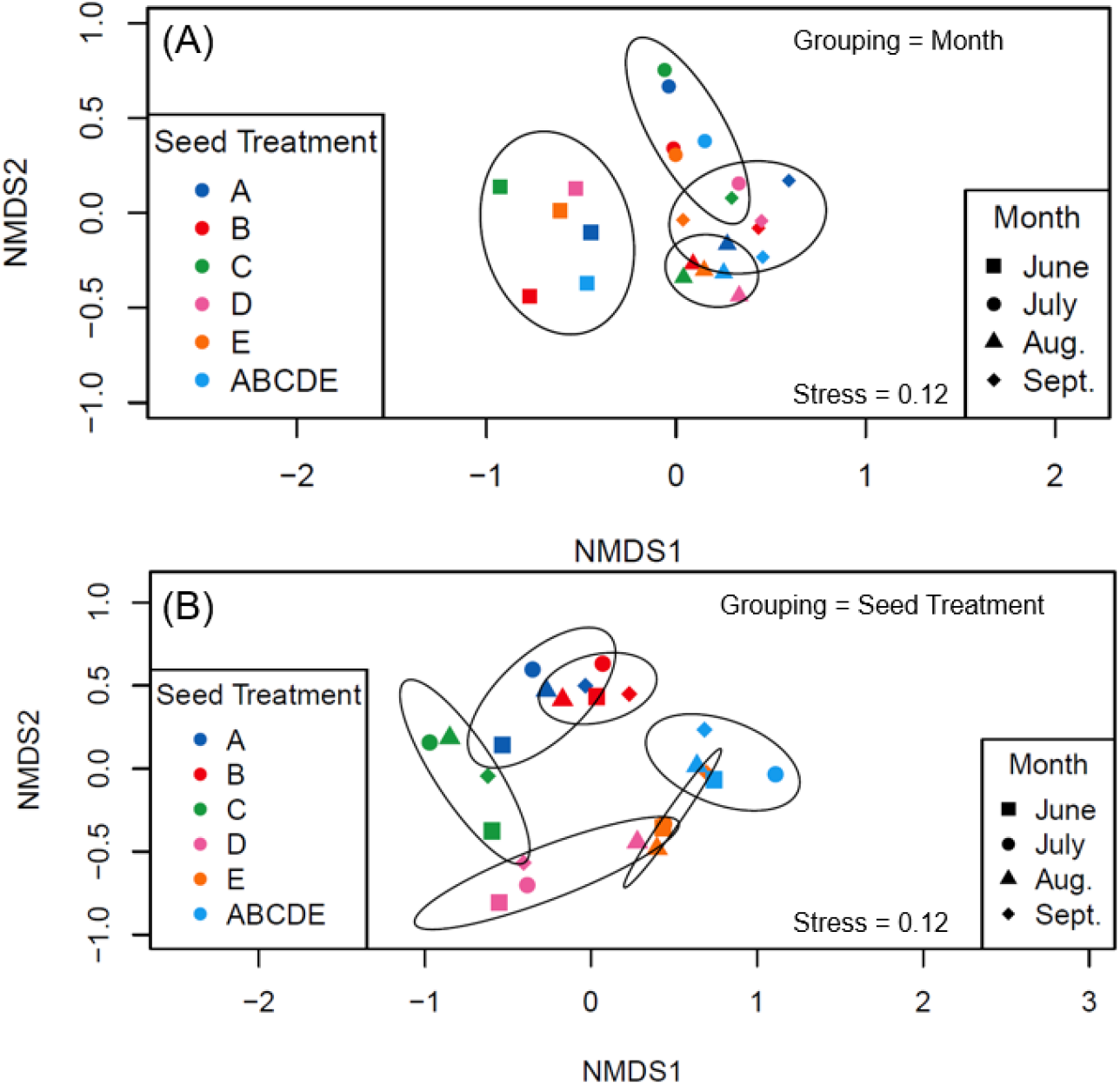
Nonmetric Multidimensional Scaling with Bray – Curtis dissimilarity graphs of the first year established communities within (A) RSC communities grouped by seed treatment and (B) ORD communities grouped by month. Seed treatment indicated by color and shapes indicate month of data collection. Ellipses are 95% confidence intervals.

Within the ORD experimental restoration site, we found no significant community-level differences between seed mix treatments. However, within our PERMANOVA of community composition based on Shannon’s Diversity Index, we observed a significant difference among our ORD communities based on month alone (pseudo-*F*= 0.385, *p*<0.001; Table 3). These results indicate that any differences in community diversity was not due to seed mix treatments but were primarily explained by the growing season (Fig. 3A). Pairwise comparisons found that plant community composition in June was significantly different from the later seasonal communities in August and September (Table S8). These results suggest that community diversity observed across the restoration site was different in June than was observed later in the season.

**Table 3.** PERMANOVA results for community composition differences within ORD experimental seed mix treatments, using Seed Treatment, Plot Replicate, Month, and the interaction between seed treatment and month as main explanatory variables.

|  | Df | SS | MS | Pseudo F | R2 | *P* |
| --- | --- | --- | --- | --- | --- | --- |
| Seed Treatment | 5 | 0.181 | 0.036 | 2.165 | 0.077 | 0.068 |
| PlotRep | 14 | 0.228 | 0.016 | 0.975 | 0.097 | 0.509 |
| Month | 3 | 0.908 | 0.303 | 18.095 | 0.385 | *0.001* |
| Bare Ground | 1 | 0.018 | 0.018 | 1.079 | 0.008 | 0.314 |
| Thatch | 1 | 0.001 | 0.001 | 0.061 | 0.000 | 0.928 |
| Treatment:Month | 15 | 0.352 | 0.023 | 1.401 | 0.149 | 0.157 |
| Residuals | 40 | 0.669 | 0.017 | 0.284 |  |  |
| Total | 79 | 2.358 | 1.000 |  |  |  |

## DISCUSSION

Current local seed sourcing approaches during restoration may not adequately incorporate within species genetic diversity needed to re-establish functional plant communities for adaptation to changing environmental conditions. Thus, establishing diversity within and between species for seed mixes will be critical to ensuring restoration success. Using seed source as a proxy to indicate increased genetic variation, we have empirically evaluated how community diversity establishes following the use of single and multiple-source seed mix treatments. There was no major effect of seed mix treatment type on increasing community diversity within the first year following restoration at two sites. However, community diversity across seed mix treatment types at this early stage following restoration was strongly influenced by spatial heterogeneity and by the growing season across the RSC restoration site, and strongly influenced by time at the ORD site. Community diversity within both sites was largely dominated by non-seeded species, with limited emergence of seeded species within the first year. These observations are consistent with previous restoration studies, which observed that non-seeded species may dominate restored environments during the first several years following restoration (Kaul & Wilsey 2020). Although no differences were observed in community diversity between our single and multiple-source seed mix treatments, our results suggest that first-year restoration communities are influenced by heterogeneity in a restoration site and temporally by the growing season. Thus, land-use history may be important in influencing plant establishment and persistence over time, particularly in the first year following restoration.

### Seed Viability

Although non-seeded species were expected in the first year, variation in seed viability within our seed mixes (ranging from 20-88% for the RSC site) may have impacted first year emergence. For seed viability testing, 7-37% of seeds were considered “dormant” and therefore may have germinated within the first year, but could emerge in subsequent years provided that environmental conditions in the future are favorable for germination. In addition, seed predation and seedling herbivory may have reduced establishment success during the first year. Herbivore disturbance can mediate non-seeded species dispersal through selective seed herbivory on native plant species (Howe & Brown 2000) which may affect overall species diversity. At the northwestern MN site, the thirteen lined-ground squirrel (*Ictidomys tridecemlineatus*) was observed, alongside nearby and within-plot gopher mounds. As our study design was aimed to mimic natural restoration practices, we did not take measures to actively exclude mammals from the restoration sites, but instead used approximately five times the standard seeding rate for each seed mix treatment type. High seeding rates are often used to mitigate potential effects of seed viability and herbivory on seedling establishment, and increase overall plant densities (Applestein et al. 2018).

### Plant Community Structure following Restoration

We compared species richness following restoration with seed mixes containing a single source per species or multiple sources per species across two restoration sites. Multi-source seed mixes were associated with greater seeded species richness at the RSC site, but not the ORD restoration site. In the first growing season following the restoration four times the number of non-seeded species were observed compared to seeded species at the ORD site, and seven times at the RSC site, respectively (Fig. 2). This is consistent with rates observed previously in grassland restoration experiments (Martin et al. 2005). Seeded species that emerged were those have evolved traits that provide competitive advantages in grassland ecosystems, such as rhizomatous root systems (Mangan et al. 2011; Dickson & Busby 2009) or mutualist fungal relationships which can promote and facilitate establishment (Busby et al. 2011). For example, *H. maximiliani* is a widespread perennial forb native to prairies in the United States and Canada (USDA). *H. maximiliani* readily established at both sites across seed treatments and is often found in remnant and restored prairies as a sub-dominant or dominant species (Dickson & Busby 2009). Previous studies have found that *H. maximiliani* is often one of the most productive forb species within plant communities as it may outcompete other species due to its rhizomatous root system that creates a spreading pattern for nutrient uptake, and thick sprouting stem that leads to increased biomass production and vegetative coverage (McKenna et al. 2019; Mangan et al. 2011; Dickson & Busby 2009). *Ratibida columnifera* was another common perennial species to establish at the ORD site and across various seed treatments. This species occurs widely throughout southern Canada, across the US Great Plains, and into Northern Mexico (USDA). In previous experiments, *R. columnifera* has been observed to have high first year survival and a life span around three years and may negatively impact the abundance of other forbs (Lauenroth & Adler 2008; Dickson & Busby 2009). The competitive advantage expressed by *R. columnifera* may be due to its establishment through a prominent taproot and strong positive relationship with arbuscular mycorrhizal fungi which aids nutrient uptake and growth (Busby et al. 2011). In addition, both species are native to our study regions, thus may exist within the seed bank currently. However, during field site visits we did not observe *H. maximiliani* at either site outside of the experimental plots. *Ratibida columnifera* was present within the RSC site but was not included in the experimental seed mixes and was not present within the plots. Evaluating what species readily establish during the early stage of a restoration may aid in future seed mix design choices to combat non-native species establishment, and to ensure early restoration success.

Both the PERMANOVA and NMDS analyses assessed plant community structure using measures of diversity from seeded and non-seeded species quantified across seed mix treatments for each site. For RSC, the seed treatment with the most diverse community established throughout the season was our multiple-source mix (ABCDE). The multi-source seed treatments were planted on the portion of the site that was once a golf course, near a remnant mesic area with surrounding woody vegetation. Several species that established solely within this treatment were persistent within the woody vegetation nearby, including *Achillea millefolium, Plantago major, P. annua*, and *Salix interior*. The presence of these species only within our multiple-source treatment plots is therefore likely influenced by the neighboring community, although as predicted this treatment had the most seeded species establish. This treatment was significantly different from all other seed mix treatments, except ‘E’ which was compositionally more similar during later seasonal months (Table S7). Given the spatial proximity of the ‘ABCDE’ and ‘E’ treatments, similar communities likely arose due to local site conditions, including below ground nutrient resources and varying seed banks across the site. Community composition at RSC also varied over time in response to seed mix treatments (Table 2). However, the spatial differences observed in community composition were maintained throughout the growing season.

Although multi-source seed mixes were associated with greater sown species richness within the RSC plots, total sown species richness was greater across all ORD plots, but not different across seed treatments (Fig. 2). The increase in total seeded species richness could indicate there was less competition from non-seeded species which may allow for increased establishment, or seeded species already existed within the soil seed banks. Although seed treatment did not appear to influence sown species establishment within ORD plots, growing season influenced communities with similar community diversity establishing throughout the growing season (Fig. 3b). Pairwise comparisons of community diversity across time indicated that June was the only month that was significantly different from the community present in later months. This may indicate that early season emergence drives the formation of community structure across time. These data provide a baseline understanding of site-specific community diversity to monitor composition change over time and across seed treatments.

Comparison across sites suggests the different patterns of diversity and those factors that structure diversity across sites are likely associated with different land-use histories. The experimental seed mix treatments at the ORD restoration site were established on an old agricultural field with active management for hay production. The site has experienced similar land-use history, which has likely largely homogenized the above and belowground plant community, currently dominated by smooth brome (*B. inermis*) and alfalfa (*Medicago sativa*). The influence of agricultural activity and dominance of smooth brome and alfafa has also likely contributed to further homogenization of the associated seed bank, reducing richness and diversity of the non-seeded species community (Bekker et al. 1997). In contrast to the homogeneity observed at the ORD site, the land-use history at RSC was more heterogeneous, which may have contributed to spatial variation in plant community establishment across the site. Interestingly, while the ORD community structure did not exhibit differences associated with seed mix treatment, the RSC site did exhibit significant differences across seed mix treatments.

Single-source seed treatments A, B, and C were established on a portion of the site that was once planted with brome and alfalfa for haying purposes. In contrast, seed treatments D, E, and the multiple-source mix ABCDE were established on a portion of the site that was a golf course up until 2015. Combined, land use history and varying impacts of the seed bank and nutrient profile across the site suggests there is substantial heterogeneity across the site that may have influenced emergence following application of seed treatments. Despite site preparation methods used to prevent non-seeded species establishing within plots these differences may be reflected in the site-level differences as opposed to seed mix application. Thus site-level differences are due to spatial heterogeneity within the soil seed bank and nutrient availability associated with land-use history impacting community establishment regardless of seed mix treatment.

Land use history can play an important role influencing how restoration communities establish over time (Cousins et al. 2009; Grman et al. 2013). Spatial heterogeneity across a restoration site could influence soil nutrient resources across the site and the associated species that may persist within the seed bank (Ricklefs 1977; Bakker et al. 2003). Where greater nutrient loading is observed, increased competition and exclusion between seeded and non-seeded species for resources could be observed (Eskelinen et al. 2021; Stotz et al. 2019). Aggressive non-seeded species often outcompete natives along nutrient load gradients leading to a subsequent loss of available soil nutrient resources. This can have substantial impacts to native plant diversity both above and belowground (Stevens & Carson 2002; Wilson & Tilman 1993; Eskelinen et al. 2021). Thus, heterogeneity in the soil nutrients or lack thereof likely impacted how communities established at both sites, but data on emergence provide a baseline to monitor how patterns in community composition may change over time.

Although we were interested in which seeded species established within our seed mix treatments, non-seeded species may also be important components to consider when evaluating these experimental communities over time. In a previous study Kaul & Wilsey (2020) noted that non-seeded weedy species abundance was the strongest predictor of species richness and diversity in grassland restorations, regardless of the age of the restoration. The most common non-seeded species to establish within our communities were introduced species, including cool-season grasses *B. inermis* and *Poa pratensis*. These species typically outcompete natives for resources, including both nutrient and light availability (reviewed in D’Antonio & Meyerson 2002). *Poa pratensis* establishes early in the spring before many native forbs, thus early establishment and the consequent increased growing season may provide a competitive advantage over native species (DeKeyser et al. 2015). *Bromus inermis* also establishes readily in the spring and is a commonly planted pasture grass that readily forms a quickly establishing monoculture through a rhizomatous root system (Stotz et al. 2019). The aggressive establishment of *B. inermis* often leads to outcompeting and displacing native species which may lead to decreased plant diversity and community homogenization of a site when it becomes an established invader (Stotz et al. 2019; DiAllesandro et al. 2013). The prevalence of these well-known invasive species within our treatments, despite our pre-seeding site prep to limit non-seeded species establishment may indicate that more work is needed to successfully limit and manage their establishment during restoration. Considering how these species establish may be critical to restoration success as it may require more effort to shift these communities back to native species (Martin & Wilsey 2014). Additionally, genetic variation within seeded species used within seed mixes may mitigate some of the negative impacts of invasives. Genetic variation may increase the diversity of genotypes that establish increasing the probabilities of producing a self-sustaining, persistent population that can evolve over generations. Evaluating which non-seeded species establish and tracking their abundance in the early stages of a restoration will help guide restoration expectations and community management practices over time.

Single versus multiple source seed mix treatments did not have an impact on community composition diversity in the first year of restoration establishment. Our results suggest that early emergence and diversity within a plant community following restoration is largely influenced by land-use history. In addition, first-year emergence following restoration may be largely insensitive to seed mix type if non-seeded species in the seedbank are able to outcompete seeded species during establishment. Previous studies have shown that first year emergence positively influences seeded species abundance and richness several years following restoration (Applestein et al. 2018; Geaumont et al. 2019) Thus, while there is some evidence to suggest seed mix type may impact the diversity of established species, long-term assessments over multiple years will be necessary to quantify the full impact of seed mix type has to community diversity and restoration success over time. Evaluating what seeded and non-seeded species establish in the first year of a restoration will help inform future restoration plans for long-term restoration success. Indeed, identifying those seeded species that may have the competitive ability to readily establish may be needed during the design of seed mixes, both identifying those species that should be included and the proportion of seed that may be necessary to maintain those species over time.

Understanding the role within and among population genetic variation has on native grassland restorations may have substantial implications to seed mix design recommendations. We assumed here that a multi-population seed mix reflects increased genetic variation, however, the degree to which population sources impact standing genetic variation within seed mixes remains to be tested. Future work should include a genetic analysis of populations in single and multi-source seed mixtures. Finally, although initial establishment results may be important to early restoration success, longer-term monitoring will be necessary to evaluate the impact seed mix treatment may have to community structure over time. Combined, genetic analysis and longer-term monitoring of seed mix treatments will provide information needed for land managers to establish seed sourcing guidelines critical to restoration in a changing environment.

## ACKNOWLEDGEMENTS

We thank the dedicated staff and volunteers at the Samuel H. Ordway Prairie Preserve and Minnesota State University Moorhead Regional Science Center and summer field crews. We also thank The Nature Conservancy, US Fish & Wildlife Service, North and South Dakota Game & Fish, South Dakota School Trust Lands, and Minnesota Department of Natural Resources for access to survey sites and seed collection. We are grateful to J. Waraniak and J. Braasch for statistical help, and to R. Hedlund for conducting vegetation surveys in South Dakota. Funding was provided by the Clean Water Land & Legacy Amendment, Doris Duke Charitable Foundation, Prairie Pothole Joint Venture, Wildlife Conservation Society’s Climate Adaptation Fund, and The Nature Conservancy. The authors declare no conflict of interest. Data accessibility will be provided.

