## Supplemental for "Seed sourcing for climate-resilient grasslands: the role of seed source diversity during early restoration establishment"

Table S1. Individual guides with links used to calculate the species-specific number of seeds found per gram for each species used in restoration seed treatments.

| **Species Scientific Name** | **Guide Source** |
| --- | --- |
| *Amorpha canescens* | Prairie Moon, https://www.prairiemoon.com/ |
| *Anemone cylindrica* | Native Seed Production Manual, https://tallgrassprairiecenter.org/sites/default/files/native_seed_production_manual.pdf |
| *Artemisia frigida* | L&H Seed, http://www.lhseeds.com/artemisia-frigida-fringed-sagebrush |
| *Bouteloua curtipendula* | Native Seed Production Manual |
| *Bouteloua gracilis* | Prairie Moon |
| *Dalea purpurea* | Native Seed Production Manual |
| *Echinacea angustifolia* | Prairie Moon |
| *Geum triflorum* | Prairie Moon |
| *Helianthus maximiliani* | Prairie Moon |
| *Helianthus pauciflorus* | Prairie Moon |
| *Hesperostipa comata* | USDA Plant Database, https://plants.usda.gov/plantguide/pdf/pg_heco26.pdf |
| *Liatris punctata* | Prairie Moon |
| *Pediomelum argophyllum* | Shirley 1994. Restoring the Tallgrass Prairie: An Illustrated Manual for Iowa and the Upper Midwest. |
| *Potentilla arguta* | Prairie Moon |
| *Ratibida columnifera* | Prairie Moon |
| *Schizachriym scoparium* | Native Seed Production Manual |
| *Solidago rigida* | Native Seed Production Manual |

Table S2. Pairwise geographic distances in kilometers between individual seed collection sites from remnant prairies sampled throughout northwestern Minnesota. Pairwise distances ranged from 3.11 km to a maximum of 215.13 km.

|  | GRP | AGD | TWI | FMB | ZIM | BIC | FLI | BLU | OLS | BLA | HAN | SEV | POM |
| --- | --- | --- | --- | --- | --- | --- | --- | --- | --- | --- | --- | --- | --- |
| AGD | 22.84 | - | - | - | - | - | - | - | - | - | - | - | - |
| TWI | 59.89 | 37.09 | - | - | - | - | - | - | - | - | - | - | - |
| FMB | 52.88 | 30.40 | 8.62 | - | - | - | - | - | - | - | - | - | - |
| ZIM | 68.45 | 46.29 | 17.86 | 25.37 | - | - | - | - | - | - | - | - | - |
| BIC | 74.88 | 52.20 | 15.43 | 22.10 | 22.36 | - | - | - | - | - | - | - | - |
| FLI | 73.66 | 50.89 | 13.81 | 21.27 | 19.25 | 3.11 | - | - | - | - | - | - | - |
| BLU | 96.73 | 74.05 | 37.09 | 43.92 | 37.37 | 21.86 | 23.31 | - | - | - | - | - | - |
| OLS | 94.64 | 71.96 | 36.44 | 44.93 | 27.39 | 25.98 | 24.95 | 19.37 | - | - | - | - | - |
| BLA | 114.37 | 91.65 | 55.66 | 63.97 | 46.88 | 43.34 | 43.10 | 26.97 | 19.74 | - | - | - | - |
| HAN | 150.04 | 127.23 | 90.52 | 98.40 | 83.16 | 76.63 | 77.13 | 55.98 | 55.85 | 36.32 | - | - | - |
| SEV | 183.54 | 161.70 | 128.18 | 136.75 | 115.42 | 117.27 | 116.70 | 100.31 | 91.93 | 74.18 | 51.64 | - | - |
| POM | 150.51 | 127.68 | 90.67 | 98.23 | 84.86 | 76.19 | 77.02 | 54.81 | 57.49 | 38.75 | 8.17 | 58.52 | - |
| STA | 215.13 | 193.04 | 158.63 | 167.11 | 146.77 | 146.80 | 146.53 | 128.29 | 122.19 | 103.47 | 74.50 | 32.36 | 79.33 |

Table S3. Geographic distances in km between individual seed collection sites to established restoration plots at the RSC site within northwestern Minnesota. Distances ranged from 2.22 km to 129.27 km.

| Site Code | Distance (km) |
| --- | --- |
| GRP | 94.82 |
| AGD | 72.10 |
| TWI | 35.08 |
| FMB | 42.06 |
| ZIM | 35.15 |
| BIC | 19.96 |
| FLI | 21.28 |
| BLU | 2.22 |
| OLS | 17.96 |
| BLA | 27.25 |
| HAN | 57.37 |
| SEV | 100.99 |
| POM | 56.43 |
| STA | 129.27 |

Table S4. Pairwise geographic distances in kilometers between individual seed collection sites from remnant prairies sampled throughout the Missouri Coteau region. Pairwise distances ranged from 2.38 km to a maximum of 311.56 km.

|  | MYR | GRO | COR | NBM | KRU | MUN | KOS | LSB | ORD | TEN | EUR | RYM | GDY | ARF | JNK | MIL |
| --- | --- | --- | --- | --- | --- | --- | --- | --- | --- | --- | --- | --- | --- | --- | --- | --- |
| GRO | 11.66 | - | - | - | - | - | - | - | - | - | - | - | - | - | - | - |
| COR | 27.66 | 16.10 | - | - | - | - | - | - | - | - | - | - | - | - | - | - |
| NBM | 24.80 | 33.56 | 46.52 | - | - | - | - | - | - | - | - | - | - | - | - | - |
| KRU | 16.98 | 18.42 | 27.62 | 19.87 | - | - | - | - | - | - | - | - | - | - | - | - |
| MUN | 27.47 | 28.09 | 33.68 | 22.02 | 10.56 | - | - | - | - | - | - | - | - | - | - | - |
| KOS | 30.54 | 28.81 | 31.28 | 28.13 | 13.87 | 6.14 | - | - | - | - | - | - | - | - | - | - |
| LSB | 66.62 | 59.90 | 50.96 | 66.86 | 52.07 | 45.09 | 39.27 | - | - | - | - | - | - | - | - | - |
| ORD | 95.95 | 92.51 | 87.37 | 86.65 | 79.18 | 69.24 | 65.41 | 39.35 | - | - | - | - | - | - | - | - |
| TEN | 89.26 | 88.22 | 86.60 | 75.45 | 72.32 | 61.79 | 59.52 | 46.50 | 19.95 | - | - | - | - | - | - | - |
| EUR | 93.81 | 95.34 | 97.05 | 75.31 | 77.67 | 67.35 | 66.95 | 64.46 | 40.31 | 20.96 | - | - | - | - | - | - |
| RYM | 143.77 | 142.45 | 139.47 | 128.96 | 126.84 | 116.31 | 113.91 | 92.81 | 53.46 | 54.52 | 56.29 | - | - | - | - | - |
| GDY | 180.80 | 180.02 | 177.46 | 164.59 | 163.95 | 153.39 | 151.33 | 130.70 | 91.38 | 91.81 | 89.73 | 38.05 | - | - | - | - |
| ARF | 192.58 | 190.04 | 185.01 | 179.39 | 175.61 | 165.24 | 162.19 | 135.25 | 97.75 | 104.06 | 107.58 | 51.29 | 30.24 | - | - | - |
| JNK | 194.96 | 192.40 | 187.34 | 181.77 | 177.99 | 167.62 | 164.57 | 137.54 | 100.10 | 106.44 | 109.91 | 53.62 | 31.51 | 2.38 | - | - |
| MIL | 242.63 | 239.37 | 233.14 | 230.44 | 225.73 | 215.50 | 212.11 | 182.48 | 146.89 | 154.99 | 159.14 | 102.85 | 75.43 | 51.56 | 49.23 | - |
| NIE | 311.56 | 306.93 | 298.69 | 301.84 | 294.97 | 285.16 | 281.13 | 247.81 | 215.93 | 226.83 | 233.32 | 177.29 | 151.75 | 126.58 | 124.38 | 76.32 |

Table S5. Geographic distances in km between individual seed collection sites to established restoration plots at the ORD site within the Missouri Coteau. Distances ranged from 3.54 km to 214.04 km.

| Site Code | Distance (km) |
| --- | --- |
| MYR | 97.61 |
| GRO | 93.80 |
| COR | 88.09 |
| NBM | 89.04 |
| KRU | 80.94 |
| MUN | 71.15 |
| KOS | 67.12 |
| LSB | 39.13 |
| ORD | 3.54 |
| TEN | 23.48 |
| EUR | 43.77 |
| RYM | 53.94 |
| GDY | 91.65 |
| ARF | 96.91 |
| JNK | 99.25 |
| MIL | 145.59 |
| NIE | 214.04 |

Table S6. Cover-class method used to quantify coverage estimates for individual species, litter cover, and bare ground soil coverage modified from Daubenmire (1959). Estimates were taken for all quadrats sampled and averaged to obtain a plot-replicate level estimates of coverage.

| Code | Estimated cover range |
| --- | --- |
| 0 | 0-4% |
| 5 | 5-9% |
| 10 | 10-19% |
| 20 | 20-29% |
| 30 | 30-39% |
| 40 | 40-49% |
| 50 | 50-59% |
| 60 | 60-69% |
| 70 | 70-79% |
| 80 | 80-89% |
| 90 | 90-94% |
| 95 | 95-99% |
| 100 | 100% |

| June | | | | |  | | July | | | | | | | | | | | |
| --- | --- | --- | --- | --- | --- | --- | --- | --- | --- | --- | --- | --- | --- | --- | --- | --- | --- | --- |
| Pairs | SS | F | R2 | *p* | |  | | Pairs | SS | F | | | R2 | | | *p* |  |  |
| A vs B | 0.62 | 4.71 | 0.54 | 0.10 | |  | | A vs B | 0.72 | 10.61 | | 0.73 | | | 0.10 | | | |
| A vs C | 0.70 | 4.16 | 0.51 | 0.10 | |  | | A vs C | 0.75 | 5.28 | | 0.57 | | | 0.10 | | | |
| A vs D | 0.71 | 3.49 | 0.47 | 0.10 | |  | | A vs D | 0.41 | 2.22 | | 0.36 | | | 0.10 | | | |
| A vs E | 0.69 | 11.82 | 0.75 | 0.10 | |  | | A vs E | 0.77 | 11.55 | | 0.74 | | | 0.10 | | | |
| A vs ABCDE | 1.25 | 10.48 | 0.64 | *0.02* | |  | | A vs ABCDE | 1.27 | 13.42 | | 0.69 | | | *0.02* | | | |
| B vs C | 0.73 | 3.26 | 0.45 | 0.10 | |  | | B vs C | 0.73 | 4.12 | | 0.51 | | | 0.10 | | | |
| B vs D | 0.68 | 2.63 | 0.40 | 0.20 | |  | | B vs D | 0.57 | 2.63 | | 0.40 | | | 0.10 | | | |
| B vs E | 0.57 | 5.02 | 0.56 | 0.10 | |  | | B vs E | 0.38 | 3.75 | | 0.48 | | | 0.10 | | | |
| C vs D | 0.14 | 0.46 | 0.10 | 0.60 | |  | | C vs D | 0.21 | 0.73 | | 0.15 | | | 0.20 | | | |
| C vs E | 0.92 | 6.16 | 0.61 | 0.10 | |  | | C vs E | 0.82 | 4.67 | | 0.54 | | | 0.10 | | | |
| D vs E | 0.91 | 4.92 | 0.55 | 0.10 | |  | | D vs E | 0.65 | 2.99 | | 0.43 | | | 0.10 | | | |
| ABCDE vs B | 1.12 | 7.18 | 0.54 | *0.02* | |  | | ABCDE vs B | 1.09 | 9.27 | | 0.61 | | | *0.02* | | | |
| ABCDE vs C | 1.13 | 6.26 | 0.51 | *0.02* | |  | | ABCDE vs C | 1.12 | 6.66 | | 0.53 | | | *0.02* | | | |
| ABCDE vs D | 1.11 | 5.43 | 0.47 | *0.02* | |  | | ABCDE vs D | 0.93 | 4.77 | | 0.44 | | | *0.02* | | | |
| ABCDE vs E | 0.82 | 7.62 | 0.56 | *0.02* | |  | | ABCDE vs E | 0.60 | 5.13 | | 0.46 | | | *0.02* | | | |
| August | | | | |  | | September | | | | | | | | | | | |
| Pairs | SS | F | R2 | *p* | |  | | Pairs | SS | | F | | | R2 | | | | *p* |
| A vs B | 0.48 | 7.87 | 0.66 | 0.10 | |  | | A vs B | 0.32 | | 3.41 | | | 0.46 | | | | 0.10 |
| A vs C | 0.51 | 4.02 | 0.50 | 0.10 | |  | | A vs C | 0.57 | | 3.25 | | | 0.45 | | | | 0.10 |
| A vs D | 0.42 | 1.82 | 0.31 | 0.10 | |  | | A vs D | 0.33 | | 1.60 | | | 0.29 | | | | 0.20 |
| A vs E | 0.30 | 3.66 | 0.48 | 0.10 | |  | | A vs E | 0.61 | | 5.59 | | | 0.58 | | | | 0.10 |
| A vs ABCDE | 0.77 | 8.68 | 0.59 | *0.02* | |  | | A vs ABCDE | 0.73 | | 5.13 | | | 0.46 | | | | *0.02* |
| B vs C | 0.64 | 4.31 | 0.52 | 0.10 | |  | | B vs C | 0.69 | | 3.67 | | | 0.48 | | | | 0.10 |
| B vs D | 0.43 | 1.72 | 0.30 | 0.10 | |  | | B vs D | 0.66 | | 2.97 | | | 0.43 | | | | 0.20 |
| B vs E | 0.56 | 5.47 | 0.58 | 0.10 | |  | | B vs E | 0.45 | | 3.71 | | | 0.48 | | | | 0.10 |
| C vs D | 0.14 | 0.44 | 0.10 | 0.90 | |  | | C vs D | 0.25 | | 0.84 | | | 0.17 | | | | 0.20 |
| C vs E | 0.38 | 2.27 | 0.36 | 0.30 | |  | | C vs E | 0.62 | | 3.05 | | | 0.43 | | | | 0.10 |
| D vs E | 0.26 | 0.95 | 0.19 | 0.40 | |  | | D vs E | 0.72 | | 3.05 | | | 0.43 | | | | 0.10 |
| ABCDE vs B | 0.84 | 8.13 | 0.58 | *0.02* | |  | | ABCDE vs B | 0.87 | | 5.80 | | | 0.49 | | | | *0.02* |
| ABCDE vs C | 0.66 | 4.51 | 0.43 | *0.02* | |  | | ABCDE vs C | 0.80 | | 3.90 | | | 0.39 | | | | *0.02* |
| ABCDE vs D | 0.44 | 2.04 | 0.25 | *0.04* | |  | | ABCDE vs D | 0.77 | | 3.42 | | | 0.36 | | | | *0.02* |
| ABCDE vs E | 0.23 | 2.00 | 0.25 | 0.07 | |  | | ABCDE vs E | 0.22 | | 1.35 | | | 0.18 | | | | 0.31 |

Table S7. RSC pairwise comparisons evaluating differences in community composition by seed treatment. Data is subset by month of data collection to account for significant PERMANOVA interaction between seed treatment and month on community composition. Within this analysis the multiple-source mix communities were significantly different from all single-source mixes. except for seed source E in August and September.

Table S8. ORD pairwise comparisons on community diversity differences between month of data collection. Community compositions were significantly different in June compared to August and September.

| Pairs | SS | F | R2 | *p* |
| --- | --- | --- | --- | --- |
| June vs July | 0.16 | 1.11 | 0.10 | 0.36 |
| June vs August | 0.27 | 2.27 | 0.19 | *0.05* |
| June vs September | 0.30 | 2.51 | 0.20 | *0.02* |
| July vs August | 0.24 | 1.96 | 0.16 | 0.11 |
| July vs September | 0.20 | 1.57 | 0.14 | 0.16 |
| August vs September | 0.08 | 0.80 | 0.07 | 0.58 |

Table S9. Species collection information for northwestern MN seed mixes sorted by species, the single-source seed mix individual species were used in (A,B,C,D,E), the location code species were sourced from (code is labeled by US state of collection, region name, and a unique three letter combination identifying site), and the location of site by latitude and longitude.

| **Species Scientific Name** | **Mix** | **Location** | **Latitude** | **Longitude** |
| --- | --- | --- | --- | --- |
| *Amoprha canescens* | A | MN-ABR-AGD | 47.51152 | -96.29388 |
| *Amoprha canescens* | B | MN-ABR-BIC | 47.0507 | -96.42639 |
| *Amoprha canescens* | C | MN-ABR-BLU | 46.85638 | -96.47015 |
| *Amoprha canescens* | D | MN-ABR-BLA | 46.68888 | -96.21477 |
| *Amoprha canescens* | E | MN-ABR-POM | 46.36548 | -96.40331 |
| *Anemone cylindrica* | A | MN-ABR-AGD | 47.51152 | -96.29388 |
| *Anemone cylindrica* | B | MN-ABR-BIC | 47.0507 | -96.42639 |
| *Anemone cylindrica* | C | MN-ABR-BLU | 46.85638 | -96.47015 |
| *Anemone cylindrica* | D | MN-ABR-BLA | 46.68888 | -96.21477 |
| *Anemone cylindrica* | E | MN-ABR-SEV | 46.10707 | -95.74233 |
| *Artemisia frigida* | A | MN-ABR-AGD | 47.51152 | -96.29388 |
| *Artemisia frigida* | B | MN-ABR-BIC | 47.0507 | -96.42639 |
| *Artemisia frigida* | C | MN-ABR-BLU | 46.85638 | -96.47015 |
| *Artemisia frigida* | D | MN-ABR-FMB | 47.24906 | -96.4069 |
| *Artemisia frigida* | E | MN-ABR-SEV | 46.10707 | -95.74233 |
| *Bouteloua curtipendula* | A | MN-ABR-AGD | 47.51152 | -96.29388 |
| *Bouteloua curtipendula* | B | MN-ABR-BIC | 47.0507 | -96.42639 |
| *Bouteloua curtipendula* | C | MN-ABR-BLU | 46.85638 | -96.47015 |
| *Bouteloua curtipendula* | D | MN-ABR-HAN | 46.3671 | -96.29711 |
| *Bouteloua curtipendula* | E | MN-ABR-POM | 46.36548 | -96.40331 |
| *Dalea purpurea* | A | MN-ABR-AGD | 47.51152 | -96.29388 |
| *Dalea purpurea* | B | MN-ABR-BIC | 47.0507 | -96.42639 |
| *Dalea purpurea* | C | MN-ABR-BLU | 46.85638 | -96.47015 |
| *Dalea purpurea* | D | MN-ABR-BLA | 46.68888 | -96.21477 |
| *Dalea purpurea* | E | MN-ABR-POM | 46.36548 | -96.40331 |
| *Echinacea angustifolia* | A | MN-ABR-TWI | 47.18041 | -96.35409 |
| *Echinacea angustifolia* | B | MN-ABR-BIC | 47.0507 | -96.42639 |
| *Echinacea angustifolia* | C | MN-ABR-BLU | 46.85638 | -96.47015 |
| *Echinacea angustifolia* | D | MN-ABR-OLS | 46.866415 | -96.21658 |
| *Echinacea angustifolia* | E | MN-ABR-STA | 45.815993 | -95.748746 |
| *Geum triflorum* | A | MN-ABR-GRP | 47.716661 | -96.278738 |
| *Geum triflorum* | B | MN-ABR-FMB | 47.24906 | -96.4069 |
| *Geum triflorum* | C | MN-ABR-BLU | 46.85638 | -96.47015 |
| *Geum triflorum* | D | MN-ABR-BLA | 46.68888 | -96.21477 |
| *Geum triflorum* | E | MN-ABR-SEV | 46.10707 | -95.74233 |
| *Helianthus maximiliani* | A | MN-ABR-TWI | 47.18041 | -96.35409 |
| *Helianthus maximiliani* | B | MN-ABR-ZIM | 47.10778 | -96.14406 |
| *Helianthus maximiliani* | C | MN-ABR-BLU | 46.85638 | -96.47015 |
| *Helianthus maximiliani* | D | MN-ABR-BLA | 46.68888 | -96.21477 |
| *Helianthus maximiliani* | E | MN-ABR-STA | 45.815993 | -95.748746 |
| *Helianthus pauciflorus* | A | MN-ABR-TWI | 47.18041 | -96.35409 |
| *Helianthus pauciflorus* | B | MN-ABR-ZIM | 47.10778 | -96.14406 |
| *Helianthus pauciflorus* | C | MN-ABR-BLU | 46.85638 | -96.47015 |
| *Helianthus pauciflorus* | D | MN-ABR-BLA | 46.68888 | -96.21477 |
| *Helianthus pauciflorus* | E | MN-ABR-STA | 45.815993 | -95.748746 |
| *Liatris punctata* | A | MN-ABR-AGD | 47.51152 | -96.29388 |
| *Liatris punctata* | B | MN-ABR-BIC | 47.0507 | -96.42639 |
| *Liatris punctata* | C | MN-ABR-BLU | 46.85638 | -96.47015 |
| *Liatris punctata* | D | MN-ABR-BLA | 46.68888 | -96.21477 |
| *Liatris punctata* | E | MN-ABR-SEV | 46.10707 | -95.74233 |
| *Pediomelum argophyllum* | A | MN-ABR-TWI | 47.18041 | -96.35409 |
| *Pediomelum argophyllum* | B | MN-ABR-ZIM | 47.10778 | -96.14406 |
| *Pediomelum argophyllum* | C | MN-ABR-BLU | 46.85638 | -96.47015 |
| *Pediomelum argophyllum* | D | MN-ABR-OLS | 46.866415 | -96.21658 |
| *Pediomelum argophyllum* | E | MN-ABR-STA | 45.815993 | -95.748746 |
| *Potentilla (Drymocallis) arguta* | A | MN-ABR-POM | 46.36548 | -96.40331 |
| *Potentilla (Drymocallis) arguta* | B | MN-ABR-BIC | 47.0507 | -96.42639 |
| *Potentilla (Drymocallis) arguta* | C | MN-ABR-BLU | 46.85638 | -96.47015 |
| *Potentilla (Drymocallis) arguta* | D | MN-ABR-BLA | 46.68888 | -96.21477 |
| *Potentilla (Drymocallis) arguta* | E | MN-ABR-STA | 45.815993 | -95.748746 |
| *Schizachriym scoparium* | A | MN-ABR-TWI | 47.18041 | -96.35409 |
| *Schizachriym scoparium* | B | MN-ABR-BIC | 47.0507 | -96.42639 |
| *Schizachriym scoparium* | C | MN-ABR-BLU | 46.85638 | -96.47015 |
| *Schizachriym scoparium* | D | MN-ABR-BLA | 46.68888 | -96.21477 |
| *Schizachriym scoparium* | E | MN-ABR-SEV | 46.10707 | -95.74233 |
| *Solidago rigida* | A | MN-ABR-FMB | 47.24906 | -96.4069 |
| *Solidago rigida* | B | MN-ABR-ZIM | 47.10778 | -96.14406 |
| *Solidago rigida* | C | MN-ABR-BLU | 46.85638 | -96.47015 |
| *Solidago rigida* | D | MN-ABR-BIC | 47.0507 | -96.42639 |
| *Solidago rigida* | E | MN-ABR-STA | 45.815993 | -95.748746 |

Table S10. Species collection information for Missouri Coteau seed mixes sorted by species, single-source seed mix individual species were used in (A,B,C,D,E), the location code species were sourced from (code is labeled by US state of collection, region name, and a unique three letter combination identifying site), and the location of site by latitude and longitude.

| **Species Scientific Name** | **Mix** | **Location** | **Latitude** | **Longitude** |
| --- | --- | --- | --- | --- |
| *Amoprha canescens* | A | ND-MOCO-NBM | 46.452496 | -99.492513 |
| *Amoprha canescens* | B | ND-MOCO-KRU | 46.42352 | -99.23698 |
| *Amoprha canescens* | C | SD-MOCO-ORD | 45.716309 | -99.127932 |
| *Amoprha canescens* | D | SD-MOCO-GDY | 44.97742 | -99.63688 |
| *Amoprha canescens* | E | SD-MOCO-JNK | 44.82523 | -99.29988 |
| *Bouteloua curtipendula* | A | ND-MOCO-COR | 46.483285 | -98.887434 |
| *Bouteloua curtipendula* | B | ND-MOCO-NBM | 46.452496 | -99.492513 |
| *Bouteloua curtipendula* | C | SD-MOCO-ORD | 45.716309 | -99.127932 |
| *Bouteloua curtipendula* | D | SD-MOCO-RYM | 45.294307 | -99.455084 |
| *Bouteloua curtipendula* | E | SD-MOCO-MIL | 44.396799 | -99.145596 |
| *Bouteloua gracilis* | A | ND-MOCO-NBM | 46.452496 | -99.492513 |
| *Bouteloua gracilis* | B | ND-MOCO-KRU | 46.42352 | -99.23698 |
| *Bouteloua gracilis* | C | SD-MOCO-ORD | 45.716309 | -99.127932 |
| *Bouteloua gracilis* | D | SD-MOCO-GDY | 44.97742 | -99.63688 |
| *Bouteloua gracilis* | E | SD-MOCO-JNK | 44.82523 | -99.29988 |
| *Dalea purpurea* | A | ND-MOCO-NBM | 46.452496 | -99.492513 |
| *Dalea purpurea* | B | ND-MOCO-KOS | 46.301838 | -99.19845 |
| *Dalea purpurea* | C | SD-MOCO-ORD | 45.716309 | -99.127932 |
| *Dalea purpurea* | D | SD-MOCO-TEN | 45.780217 | -99.367829 |
| *Dalea purpurea* | E | SD-MOCO-JNK | 44.82523 | -99.29988 |
| *Echinacea angustifolia* | A | ND-MOCO-GRO | 46.54656 | -99.07637 |
| *Echinacea angustifolia* | B | ND-MOCO-LSB | 46.025487 | -98.88189 |
| *Echinacea angustifolia* | C | SD-MOCO-ORD | 45.716309 | -99.127932 |
| *Echinacea angustifolia* | D | SD-MOCO-GDY | 44.97742 | -99.63688 |
| *Echinacea angustifolia* | E | SD-MOCO-MIL | 44.396799 | -99.145596 |
| *Geum triflorum* | A | ND-MOCO-MYR | 46.575713 | -99.222739 |
| *Geum triflorum* | B | ND-MOCO-KRU | 46.42352 | -99.23698 |
| *Geum triflorum* | C | ND-MOCO-GRO | 46.54656 | -99.07637 |
| *Geum triflorum* | D | ND-MOCO-COR | 46.483285 | -98.887434 |
| *Geum triflorum* | E | ND-MOCO-MUN | 46.3308 | -99.26634 |
| *Helianthus maximiliani* | A | ND-MOCO-NBM | 46.452496 | -99.492513 |
| *Helianthus maximiliani* | B | ND-MOCO-KRU | 46.42352 | -99.23698 |
| *Helianthus maximiliani* | C | SD-MOCO-ORD | 45.716309 | -99.127932 |
| *Helianthus maximiliani* | D | ND-MOCO-COR | 46.483285 | -98.887434 |
| *Helianthus maximiliani* | E | SD-MOCO-MIL | 44.396799 | -99.145596 |
| *Helianthus pauciflorus* | A | ND-MOCO-GRO | 46.54656 | -99.07637 |
| *Helianthus pauciflorus* | B | ND-MOCO-LSB | 46.025487 | -98.88189 |
| *Helianthus pauciflorus* | C | SD-MOCO-ORD | 45.716309 | -99.127932 |
| *Helianthus pauciflorus* | D | ND-MOCO-COR | 46.483285 | -98.887434 |
| *Helianthus pauciflorus* | E | SD-MOCO-MIL | 44.396799 | -99.145596 |
| *Hesperostipa comata* | A | ND-MOCO-KRU | 46.42352 | -99.23698 |
| *Hesperostipa comata* | B | ND-MOCO-KOS | 46.301838 | -99.19845 |
| *Hesperostipa comata* | C | SD-MOCO-ORD | 45.716309 | -99.127932 |
| *Hesperostipa comata* | D | SD-MOCO-GDY | 44.97742 | -99.63688 |
| *Hesperostipa comata* | E | SD-MOCO-JNK | 44.82523 | -99.29988 |
| *Liatris punctata* | A | ND-MOCO-NBM | 46.452496 | -99.492513 |
| *Liatris punctata* | B | ND-MOCO-KRU | 46.42352 | -99.23698 |
| *Liatris punctata* | C | SD-MOCO-ORD | 45.716309 | -99.127932 |
| *Liatris punctata* | D | ND-MOCO-COR | 46.483285 | -98.887434 |
| *Liatris punctata* | E | SD-MOCO-MIL | 44.396799 | -99.145596 |
| *Pediomelum argophyllum* | A | ND-MOCO-NBM | 46.452496 | -99.492513 |
| *Pediomelum argophyllum* | B | ND-MOCO-KRU | 46.42352 | -99.23698 |
| *Pediomelum argophyllum* | C | SD-MOCO-ORD | 45.716309 | -99.127932 |
| *Pediomelum argophyllum* | D | SD-MOCO-GDY | 44.97742 | -99.63688 |
| *Pediomelum argophyllum* | E | SD-MOCO-MIL | 44.396799 | -99.145596 |
| *Potentilla (Drymocallis) arguta* | A | ND-MOCO-NBM | 46.452496 | -99.492513 |
| *Potentilla (Drymocallis) arguta* | B | SD-MOCO-JNK | 44.82523 | -99.29988 |
| *Potentilla (Drymocallis) arguta* | C | SD-MOCO-ORD | 45.716309 | -99.127932 |
| *Potentilla (Drymocallis) arguta* | D | SD-MOCO-GDY | 44.97742 | -99.63688 |
| *Potentilla (Drymocallis) arguta* | E | SD-MOCO-MIL | 44.396799 | -99.145596 |
| *Ratibida columnifera* | A | ND-MOCO-NBM | 46.452496 | -99.492513 |
| *Ratibida columnifera* | B | SD-MOCO-JNK | 44.82523 | -99.29988 |
| *Ratibida columnifera* | C | SD-MOCO-ORD | 45.716309 | -99.127932 |
| *Ratibida columnifera* | D | SD-MOCO-MIL | 44.396799 | -99.145596 |
| *Ratibida columnifera* | E | SD-MOCO-NIE | 43.804709 | -98.664164 |
| *Solidago rigida* | A | ND-MOCO-NBM | 46.452496 | -99.492513 |
| *Solidago rigida* | B | ND-MOCO-KRU | 46.42352 | -99.23698 |
| *Solidago rigida* | C | SD-MOCO-ORD | 45.716309 | -99.127932 |
| *Solidago rigida* | D | ND-MOCO-COR | 46.483285 | -98.887434 |
| *Solidago rigida* | E | SD-MOCO-MIL | 44.396799 | -99.145596 |
